# Regulation of histone methylation by automethylation of PRC2

**DOI:** 10.1101/343020

**Authors:** Xueyin Wang, Yicheng Long, Richard D. Paucek, Anne R. Gooding, Thomas Lee, Thomas R. Cech

## Abstract

Polycomb Repressive Complex 2 (PRC2) is a histone methyltransferase whose function is critical for regulating transcriptional repression in many eukaryotes including humans. Its catalytic moiety EZH2 is responsible for the tri-methylation of H3K27 and also undergoes automethylation. Using mass spectroscopic analysis of recombinant human PRC2, we identified three methylated lysine residues (K510, K514, K515) on a disordered but highly conserved loop of EZH2. These lysines were mostly mono- and di-methylated. Either mutation of these lysines or their methylation increases PRC2 histone methyltransferase activity. In addition, mutation of these three lysines in HEK293T cells using CRISPR genome-editing increases global H3K27 methylation levels. EZH2 automethylation occurs intramolecularly (*in cis*) by methylation of a pseudosubstrate sequence on the flexible loop. This post-translational modification and *cis*-regulation of PRC2 are analogous to the activation of many protein kinases by autophosphorylation. We therefore propose that EZH2 automethylation provides a way for PRC2 to modulate its histone methyltransferase activity by sensing histone H3 tails, SAM concentration, and perhaps other effectors.

## INTRODUCTION

Lysine methylation is tightly associated with the regulation of gene expression and epigenetic inheritance. A group of enzymes called methyltransferases use S-adenosyl-L-methionine (SAM) as methyl donors and catalyze the transfer of methyl groups to the ε-amino group of lysine side chains. One major group of methyltransferases contains a catalytic SET domain. The SET domain is folded into a β-sheet structure, and a catalytic tyrosine residue at the center is paramount for the transfer of the methyl group^1^.

One prominent lysine methyltransferase is Polycomb Repressive Complex 2 (PRC2), which is the sole enzymatic complex capable of catalyzing deposition of methyl groups onto lysine 27 of histone H3. PRC2 participates in the repression of genes in mammalian cells in processes such as cellular differentiation and embryonic development. Recently it was discovered that PRC2 regulates transcription by methylating non-histone targets as well.^2^

From a surge of findings in the last decade, PRC2 has been suggested to be involved in a number of disease processes, including multiple types of cancer, cardiac hypertrophy, Huntington’s disease, and latency of viral infections including HIV and HSV^3–5^. Accordingly, understanding how PRC2 is regulated holds substantial medical potential. The regulation of PRC2 has so far been known to occur through the recruitment of various accessory proteins and binding of RNA^6,7^. For instance, the accessory protein JARID2 has been shown to substantially increase PRC2 enzymatic activity^8^. Furthermore, RNA molecules containing short stretches of guanines bind to PRC2^9^ and inhibit its HMTase activity^10,11^ by inhibiting PRC2 binding to nucleosomes^12,13^, more specifically the linker regions of nucleosomes^13^.

However, other points of regulation are likely to exist, because PRC2 is known to undergo a variety of covalent post-translation modifications including phosphorylation, sumoylation, and methylation^8,14^. Indeed, PRC2 has long been thought to auto-methylate^2,8,13,15,16^. However, a role for this automethylation has not yet been described.

In the present study, we interrogated PRC2 automethylation and addressed its mechanistic and functional importance. Using biochemical and mass spectroscopic (MS) approaches, we found that PRC2 is automethylated at three lysines on a novel and evolutionarily conserved flexible loop in EZH2. Remarkably, methylation of this loop was found to substantially stimulate PRC2-catalyzed H3K27 methylation. We also found that PRC2 automethylation occurs *in cis*. Taken together, our data reveal that automethylation of the methylation loop in EZH2 leads to PRC2 stimulation, promoting deposition of histone methylation marks. This study suggests a regulatory role of PRC2 automethylation in modulating its histone methyltransferase activity in response to H3 and SAM concentration and perhaps other effectors.

## RESULTS

### Human PRC2 is methylated on the EZH2 component

PRC2 automethylation occurs during histone methyltransferase (HMTase) assays. Automethylation appeared to occur on the EZH2 and/or SUZ12 subunits (**Figure 1A**), which have similar molecular weights and therefore run as one overlapping band on SDS-PAGE. To unambiguously identify which subunit is methylated, we utilized PRC2 complexes in which a single subunit was MBP-tagged and therefore had retarded electrophoretic mobility. As shown in **Figure 1B**, EZH2, the catalytic subunit, is the main target of automethylation.

**Figure 1.**
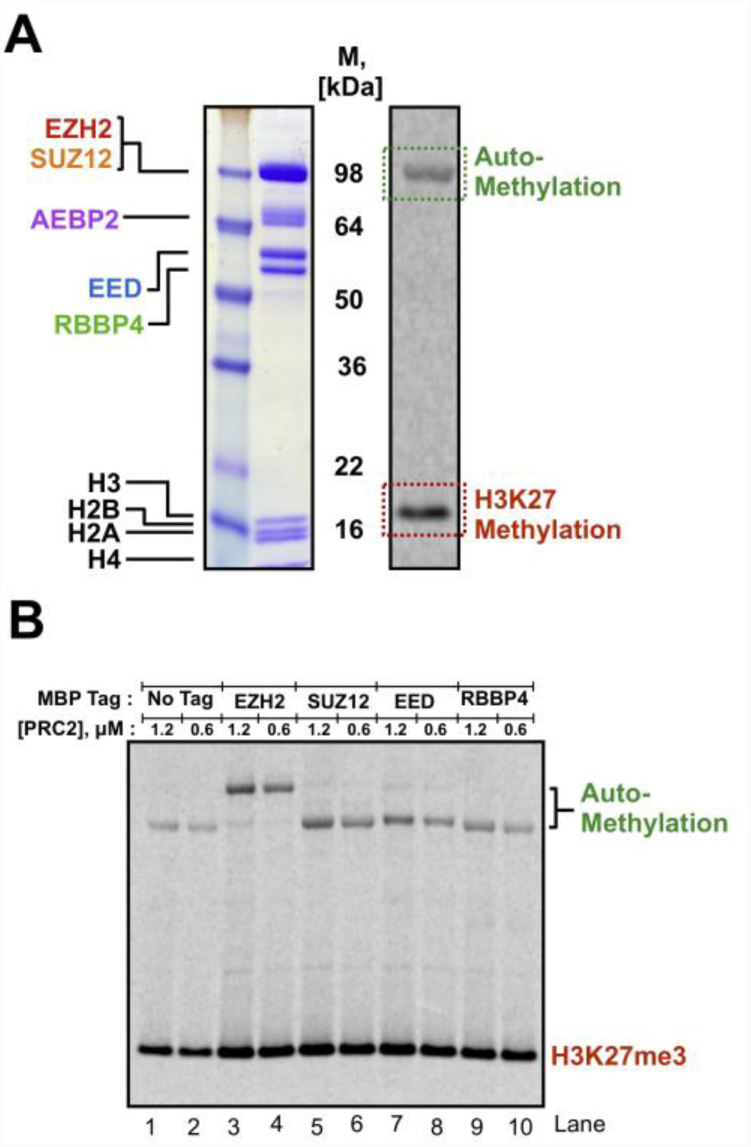
The EZH2 component of PRC2 is methylated. **A.** HMTase assays showing PRC2 enzymatic activity to mononucleosome and automethylation of PRC2. **B.** HMTase assays showing EZH2 is the subunit methylated by PRC2. HMTase assays were performed with one subunit containing an uncleavable MBP tag and mononucleosome, in the presence of cofactor ^14^C-SAM.

### Automethylation occurs at three sites on a conserved flexible loop of EZH2

To determine which EZH2 amino acid residues were being methylated, we incubated PRC2 with 10 mM SAM (unlabeled) under standard HMTase assay conditions, then subjected the protein mixtures to rare-cutting protease digestion, and lastly analyzed the samples using mass spectrometry (**Figure 2**). To avoid potential peptide bias introduced by protease digestion, independent MS experiments were performed using either Arg-C or Chymotrypsin as protease.

**Figure 2.**
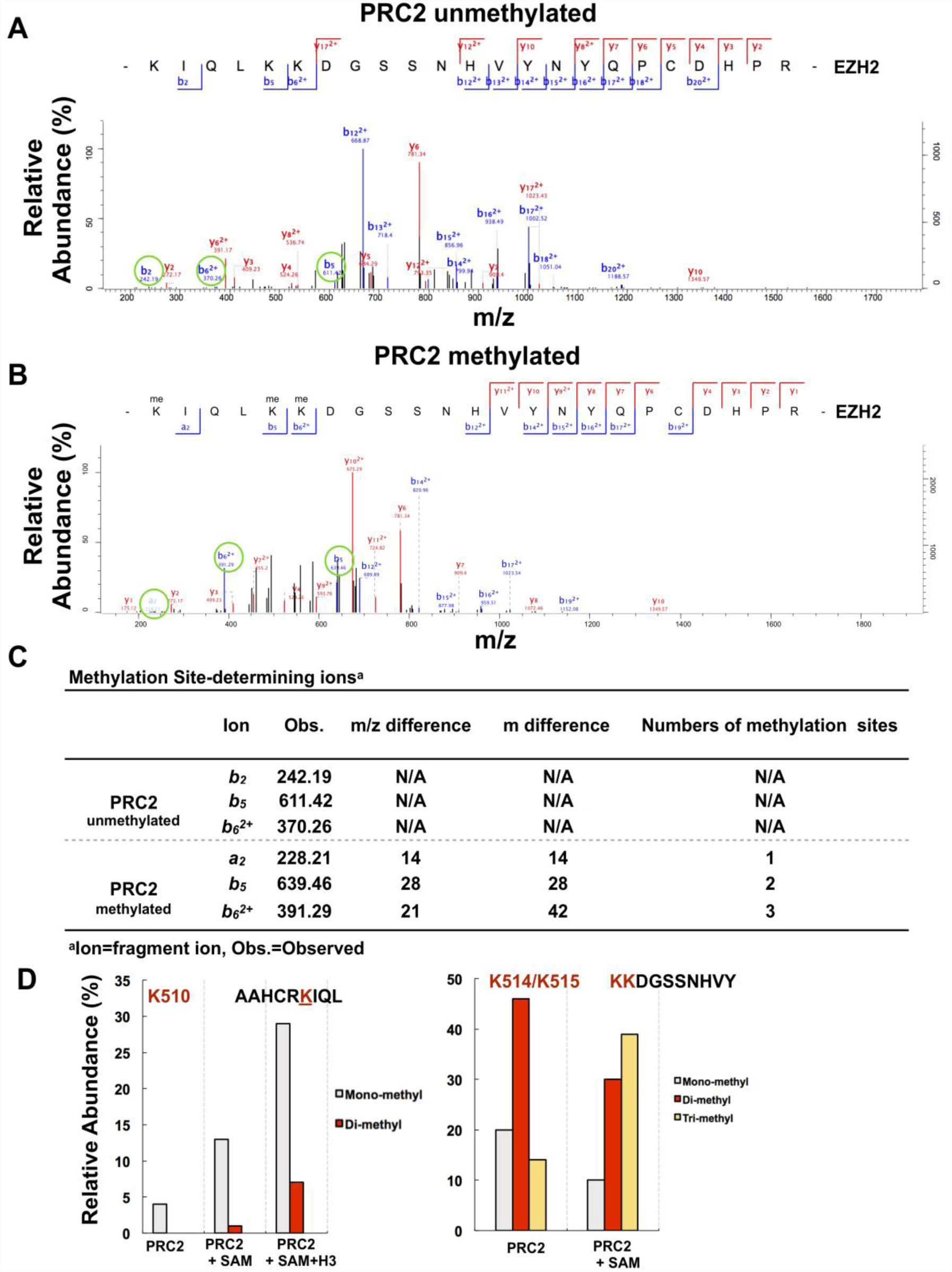
Identification of the key residues automethylated by PRC2. **A.** Mass spec analysis of unmethylated PRC2. **B.** Mass spec analysis of methylated PRC2 showing K510, K514 and K515 residues are methylated during HMTase assays. **C.** Methylation site-determining ions, circled in green in panels A and B, indicate that the indicated fragments are mono-, di- and tri-methylated. **D.** Relative abundance of fragment ions at the K510 site (left). Relative abundance of methylated isoforms of a peptide that contains K514 and K515 (right).

The additional mass due to a single methyl modification is 14 AMU. By comparing peptide masses between an unmethylated PRC2 sample (**Figure 2A**) and a methylated PRC2 sample (**Figure 2B**), automethylation marks were mapped to three lysine residues in EZH2 that exist in close proximity to one another: K510, K514, and K515 (**Figure 2C**). The expected theoretical masses of methyl-modified peptides were in good agreement with experimental observations (**Figure 2C**). Importantly, MS experiments using either Arg-C or Chymotrypin identified the same methylation sites.

Automethylation was increased in the presence of substrate H3. Three independent MS experiments were performed on samples of either PRC2, PRC2 + SAM, or PRC2 + SAM + H3. K510 was mostly mono-methylated, as illustrated in **Figure 2D** (left). The data revealed: (1) A fraction of PRC2 (7%) was already automethylated at K510 in the recombinant protein purified from insect cells; (2) The incubation of SAM with PRC2 *in vitro* increased the abundance of mono- and di-methylated peptides (from 7% to 20%); (3) The addition of H3 to a mixture of SAM and PRC2 further increased the abundance of mono-, and di-methylated peptides (from 20% to 50%). The third finding might suggest that upon binding to H3, PRC2 may undergo a conformational change that favors the automethylation of EZH2.

Because K514 and K515 are adjacent, it has been difficult to determine their methylation distribution. For example, the chymotryptic peptide KKDGSSNHVY was observed to be tri-methylated (**Figure 2D**, right), but we cannot distinguish K(me2)K(me)DGSSNHVY from K(me)K(me2)DGSSNHVY, and trimethylation of K514 or K515 would also result in the same m/z for the peptide. Other PTMs^14^ reported to decorate PRC2, such as phosphorylation and sumoylation, were not found in our MS analysis of recombinant PRC2 expressed in insect cells.

The three methylation sites (K510, K514, K515) exist on a disordered loop of EZH2 (i.e., not seen in the crystal structures^17,^^18^, nor in the cryo-EM reconstructions of PDB: 6C23 and 6C24^19^). This disordered loop in EZH2 (hereafter referred to as the “methylation loop”) extends from position 474 at the end of the SANT2 domain to position 528 at the beginning of the CXC domain (**Figure 3A**). The methylation loop shows striking sequence conservation not only between human and other mammalian homologs, but also with *Drosophila melanogaster* (**Figure 3B**). Notably, the three automethylation sites are well conserved.

**Figure 3.**
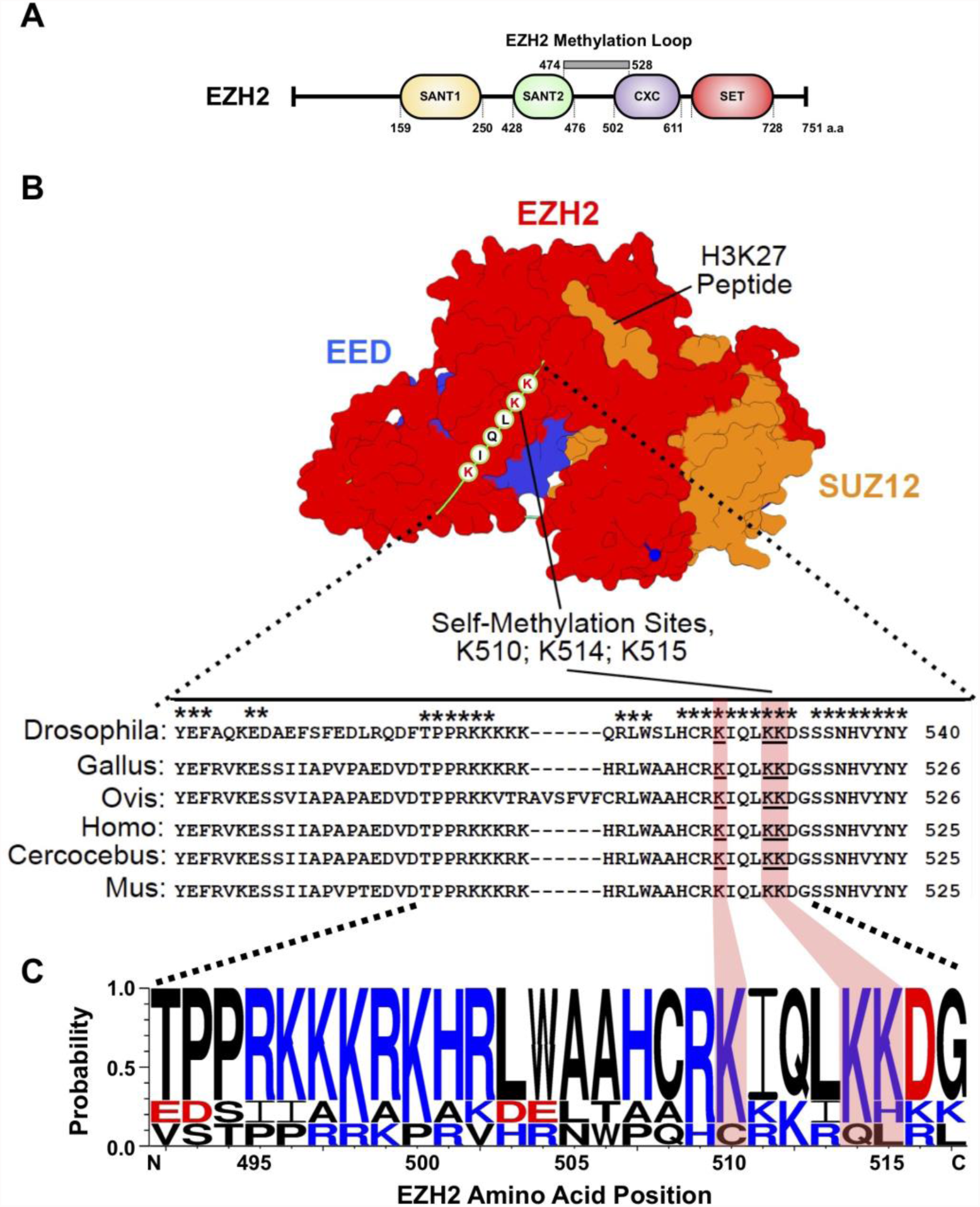
Key methylated residues in PRC2 map to a flexible and conserved charged loop in EZH2. **A.** The EZH2 methylation loop overlaps the regions flanking the SANT2 and CXC domains. **B.** The three methylated lysines (highlighted) are conserved as shown in sequence alignment. Surface representation of crystal structure of a PRC2 subcomplex from PDB: 5HYN. **C.** Sequence alignments of species shown in B show extensive conservation of a basic motif in EZH2. Blue amino acids indicate basic residues; red amino acids indicate charged residues. Methylation sites at K510, K514, and K515 are highlighted.

Another noteworthy property of the methylation loop is the large cluster of positive charges. This is illustrated in **Figure 3C** by the sequence logo representation of a selected region (residues 490-520) of the methylation loop, where the blue letters indicate positively charged residues. Given the stunning phylogenetic conservation of the methylation sites and charged residues in the EZH2 methylation loop, we hypothesized that this region may serve regulatory roles analogous to disordered loops seen in many protein kinases; phosphorylation causes a conformational change of the loop that allows substrate to bind^20^. The regulatory role of this disordered region of EZH2 (489-494) by interacting with RNA has also been recently demonstrated ^21^.

### EZH2 methylation occurs *in cis*

To better understand how the methylation loop may regulate EZH2 enzymatic activity, we asked whether automethylation occurred by a *cis*-acting mechanism (i.e., PRC2 methylating the EZH2 loop on the same protein complex) or a *trans*acting mechanism (i.e., PRC2 methylating the EZH2 loop on a neighboring protein complex).

To distinguish these possibilities, we developed a biochemical scheme that involves performing an HMTase assay on a 1:1 mixture of active PRC2 with an MBP-tag on EZH2 (“MBP-EZH2”) and untagged PRC2 with a catalytically dead EZH2 (“dEZH2”). The MBP tag on the active complex allows the unambiguous separation of active and inactive EZH2 proteins. To generate dEZH2, we introduced a Y>F single-amino-acid mutation at position 726. The design was based on the crystal structure of the EZH2 SET domain^22^, which shows the proximity of Y726 to the H3K27 substrate and the methyl donor cofactor (**Figure 4A**); the mutation of the tyrosine prevents formation of an intermediate in the methyltransferase reaction. Following expression and purification, size-exclusion chromatography of PRC2-dEZH2 showed a chromatogram identical to WT complexes, indicating that PRC2-dEZH2 was assembled and unaggregated. As shown by the HMTase assay in **Figure 4B** and **Supplementary Figure 1A**, our dEZH2 (Y726F) variant was indeed catalytically dead and did not methylate either H3 or its methylation loop. This result also eliminated the perhaps unlikely possibility that PRC2 methylation might be catalyzed by some trace contaminant enzyme that copurified with PRC2, instead of being catalyzed by PRC2 itself.

**Figure 4.**
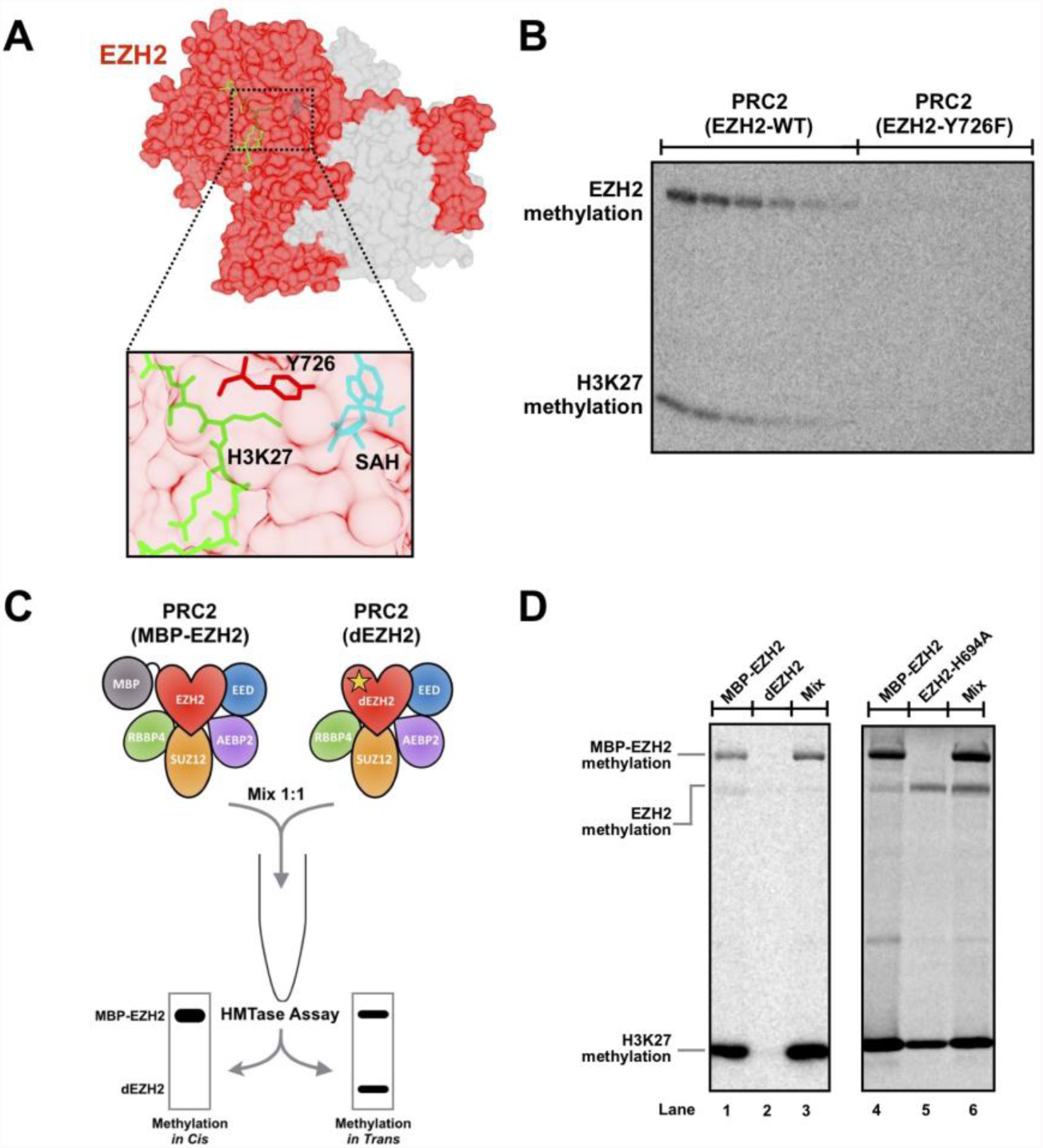
PRC2 automethylation occurs intramolecularly. **A.** PRC2 core complex crystal structure (EZH2, shown in red; EED and SUZ12, shown in gray; PDB accession number: 5HYN ^17^). Inset shows the proximity of SAH (cyan), H3 substrate (green), and the catalytic residue Y726 (red). PDB accession number: 5HYN. **B.** Missense mutation (Y726F) resulted in a catalytic dead EZH2 (dEZH2) that abolished H3K27 methylation as well as EZH2 automethylation. Reactions were carried out between PRC2 and H3, in the presence of cofactor ^14^C-SAM. **C.** Design of mixing experiments to distinguish by *cis*- and *trans*- autocatalytic reactions. **D.** Mixing experiments between WT EZH2 and dEZH2 showing that automethylation occurs *in cis*. (Left) Experiments performed by mixing WT PRC2 1:1 with PRC2 catalytic dead mutant (Y726F). (Right) Experiments performed by mixing WT PRC2 1:1 with catalytically impaired complex containing EZH2-H694A.

As shown in **Figure 4C**, considering a *cis*-pathway, one would anticipate that mixing MBP-EZH2 and dEZH2 would produce only a single methylated band corresponding to MBP-EZH2. This is expected because MBP-EZH2 would be able to methylate only itself and dEZH2 could not autocatalyze. Considering a *trans*-pathway, one would expect to observe two methylated products, because both MBP-EZH2 and dEZH2 have intact methylation loops that would be subject to methylation by MBP-EZH2. In the key experiment (**Figure 4D,** left-hand gel, Lane 3 and **Supplementary Figure 1B**), mixing of MBP-EZH2 and dEZH2 resulted in only one methylated band corresponding to MBP-EZH2, thereby confirming a *cis*-autocatalytic mechanism. The right-hand autoradiograph in **Figure 4D** shows a similar mixing experiment using a catalytically compromised H694A variant reported in the literature,^23^ which still retained partial activity under our reaction conditions. The presence of the active PRC2 in the mixture failed to restore full methylation of this mutant PRC2, again supporting methylation *in cis*.

### Mutation or methylation of the EZH2 methylation loop increases H3K27 methylation

To assess the functional importance of automethylation, we sought to produce separation-of-function PRC2 variants that preserved HMTase activity but abolished automethylation activity. Therefore, we purified a PRC2 complex with mutations (K>A) at sites 510, 515, and 515 of EZH2 (hereafter denoted as PRC2 methylmutant) and performed HMTase assays. As shown in the dashed-green box in **Figure 5A** and **Supplementary Figure 2A**, PRC2 methylmutant showed a striking reduction in automethylation signal (**Figure 5B**). The corresponding H3K27 methylation signal for the methylmutant was elevated relative to wild-type PRC2 (**Figure 5A**, dashed-red box). Indeed, quantitative analysis showed a 2-fold increase in HMTase activity (**Figure 5C**). This increase in HMTase activity for the PRC2 methylmutant suggested that EZH2 automethylation (i.e., K510^me^, K514^me^, K515^me^) might compete with H3K27 methylation. In other words, the methylation loop might be acting as a competitive inhibitor, and its mutation might remove the inhibition.

**Figure 5.**
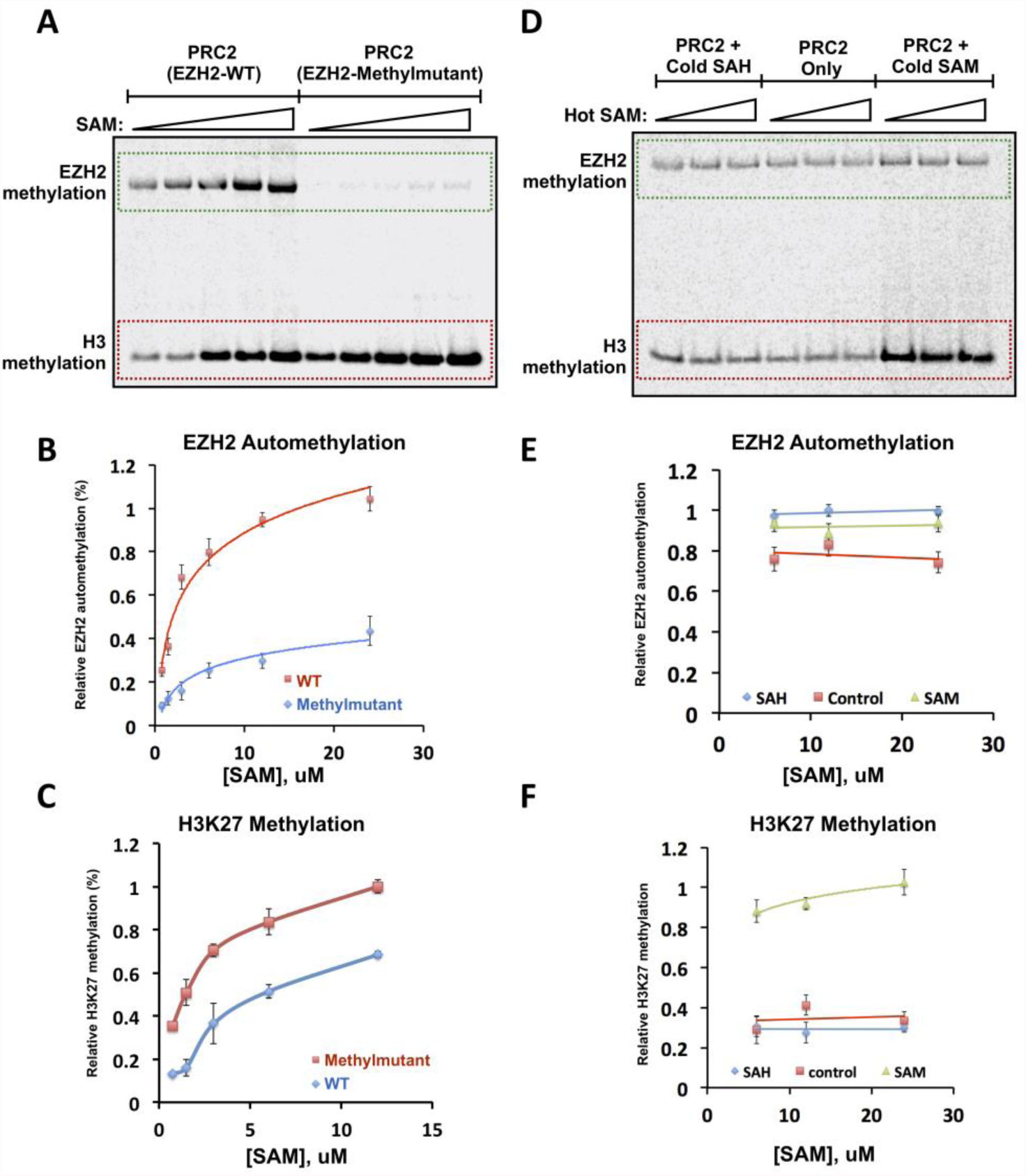
Automethylation of PRC2 activates histone substrate methylation. **A.** HMTase assays showing that triple lysine mutant has substantially reduced EZH2 automethylation but enhanced HMTase activity. **B.** Quantification of EZH2 automethylation signal (n=3) in A. **C.** Quantification (n=3) of H3K27 methylation signal in A. **D.** Pre-methylation of EZH2 enhances H3K27 methylation activity. **E.** Quantification (n=3) of EZH2 automethylation signal in D. **F.** Quantification (n=3) of H3K27 methylation signal in D.

We therefore tested the hypothesis that methylated PRC2 complexes would show attenuated HMTase activity relative to unmodified PRC2 due to competitive inhibition by the methylated loop. We proceeded by incubating PRC2 with either SAM or SAH or in the absence of cofactor. Incubation of the protein with SAM led to EZH2 automethylation (**Supplementary Figure 2B** and also confirmed by MS). Incubation of protein with SAH, or in the absence of cofactor, would leave PRC2 in an unmethylated state. HMTase assays were then performed using these pre-incubated PRC2 samples. As shown in **Figure 5D, 5E** and **Supplementary Figure 2C**, automethylation signal across the three conditions was similar. The presence of signal for the PRC2 + SAM condition simply suggests that pre-incubation did not exhaustively methylate the entire sample. Quite unexpectedly, we observed that the pre-methylation of PRC2 improved its HMTase activity (**Figure 5D and 5F**), negating our original hypothesis.

To summarize this section, we observed that (1) Mutations preventing PRC2 automethylation stimulate HMTase activity, and (2) Pre-existing automethylation stimulates HMTase activity. Thus, the mutation or methylation of sites in the EZH2 loop (i.e., K510, K514, K515) has the same consequence mechanistically. We propose that this consequence is the expulsion of the loop from the EZH2 active site, thereby freeing up the active site for HMTase activity. Thus, the flexible methylation loop of EZH2 can bind into the active site of the same EZH2 molecule, occluding entry of the H3 tail. This binding can occur in any of three productive registers, and if SAM concentrations are sufficient then the loop will be methylated and released from the active site, restoring full PRC2 activity (**Figure 6**).

**Figure 6.**
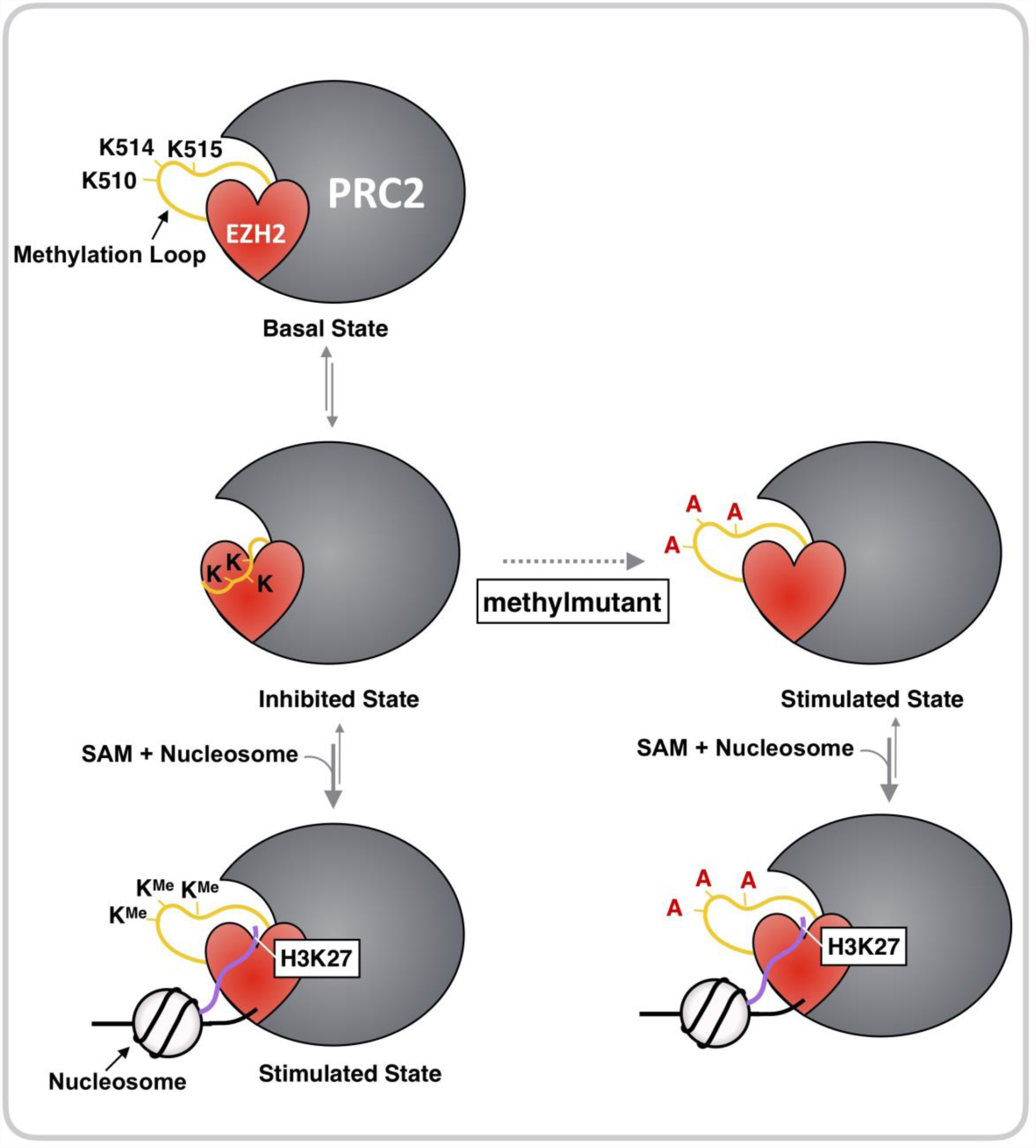
Model for autoregulation of PRC2 by the methylation loop in EZH2.

### Mutation of the EZH2 methylation loop increases global H3K27 methylation in CRISPR-edited HEK293T cells

To study the function of the EZH2 methylation loop in living human cells, we used CRISPR-Cas9 genome editing to introduce K510A, K514A and K515A mutations at the endogenous *EZH2* locus. As shown in **Figure 7A**, we performed gene editing by inserting an *EZH2* cDNA containing either the WT or mutant sequences behind the ATG start codon in exon 2. PCR followed by sequencing validation of the edited cell lines indicates that one allele is correctly edited (with WT or mutant cDNA inserted), and the other allele has in-dels right at the cleavage site (which causes frame shift and early termination of the unedited allele). These results were confirmed using Sanger sequencing (**Supplementary Figure 3**).

**Figure 7.**
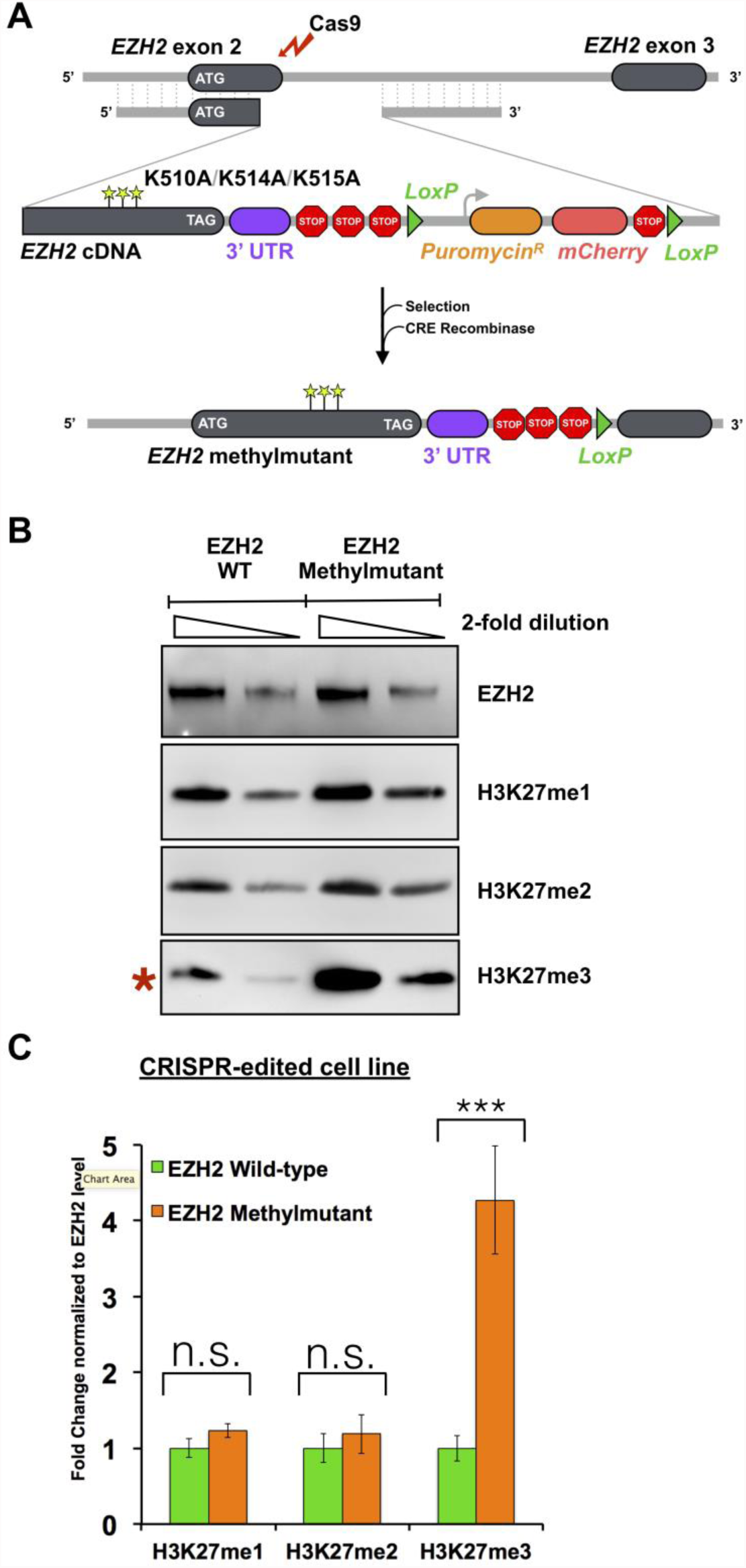
PRC2 methylmutant increases global H3K27me3 levels in HEK293T cells. **A.** CRISPR-Cas9 gene editing scheme. cDNA encoding the remaining amino acids of full length EZH2 is inserted into exon 2 of the *EZH2* locus. The cDNA to make the methylmutant encodes K>A mutations in the methylation loop at sites 510, 514, and 515. **B.** Western blot analysis of EZH2 wild-type and EZH2 methylmutant cell lines show that H3K27me3 is increased in the methylmutant cell line **C.** Quantitation of B, normalization to EZH2 levels (n=3). n.s., difference is not significant. ***P<0.001

Comparing these CRISPR-edited cell lines, we found that the expression levels of endogenous methylmutant and wild-type EZH2 were similar (**Figure 7B**). From our *in vitro* studies showing that PRC2 methylmutant has increased HMTase activity, we hypothesized that the EZH2 methylmutant cell line would show H3K27 hypermethylation. Therefore, we assayed global H3K27 methylation levels (mono-, di-, and tri-), which provide a measure of the HMTase activity of PRC2. We observed increased H3K27 tri-methylation levels in the EZH2 methylmutant cell line relative to the control cell line, while mono- and di-methylation were not affected (**Figure 7B**). As shown by the quantitative analysis in **Figure 7C**, the methylmutant increased global H3K27me3 levels by ~4 fold (4.3 ± 0.7, mean ± SD, n=3). Thus, these findings are consistent with our *in vitro* observations in **Figure 5A-C**, suggesting that the K510/K514/K515 automethylation residues serve a modulatory role in living cells.

## DISCUSSION

Autophosphorylation is a common and highly important PTM for proteins, especially those involved in cell signaling, and has been intensively studied. In contrast, only a handful of reports address protein automethylation^26,27^, even though methylation of non-histone proteins is a major PTM^23,24^. Thus, the automethylation of PRC2 studied here may be of some general interest.

A number of laboratories have observed automethylation of the core PRC2 complex^2,8,13,15,16^, attributed to its EZH2 subunit, yet basic questions pertaining to this activity have gone unanswered. What is the site of this methylation? And perhaps of more significance, what is its physiological importance? By conducting biochemical and proteomic analyses of recombinant human PRC2 complexes, we identified a conserved methylation loop in EZH2 that is modified at three lysine residues (K510, K514, and K515) via a *cis*-acting mechanism. Our data support the notion that the EZH2 methylation loop serves an autoregulatory role, and when methylated, it enhances EZH2 histone methylation activity. Furthermore, our CRISPR genome-editing of the endogenous *EZH2* gene in human cells shows the effect of these three lysines on histone methylation, but does not directly show that they function through automethylation; we have been unable to isolate enough endogenous EZH2 to confirm its methylation *in vivo*.

How does the methylation loop modulate deposition of H3K27 methyl marks? Our biochemical data and sequence comparisons best support a model in which this disordered loop acts as a pseudosubstrate for the EZH2 catalytic site (**Figure 6**). The methylation loop occupies the lysine access channel in the SET domain of EZH2 via a trio of lysine residues and prevents or slows turnover. By an intramolecular reaction, PRC2 transfers methyl groups from SAM to itself at the three possible lysines. Methylation dislodges the loop, allowing for stimulated H3 tail binding and methylation. Given the lack of a charge difference between methylated and unmethylated lysine residues, loop displacement is not driven by charge neutralization, but instead by steric effects. The Muir laboratory determined that the EZH2 active site binds strongly to linear side chains and shows little tolerance for extra steric bulk or polar groups^24^. Thus, methylation of the loop promotes release from the EZH2 active site.

How does the methylation loop sequence compare to known PRC2 methylation targets? Intriguingly, comparison of the EZH2 methylation sites identified here with sequences predicted to serve as efficient substrates for PRC2^2,23^ revealed notable insights. The Kingston laboratory recently determined that substrate regions critical for productive interaction with the PRC2 catalytic pocket typically contain an (R/K)K amino acid motif.^2^ Neighboring the targeted lysine at position −1, the arginine (R) or lysine (K) is thought to be critical due to hydrogen bonding that stabilizes peptide binding. Positions −1 of K510 and K515 in the EZH2 methylation loop are occupied by arginine (R509) and lysine (K514), respectively. Another protein target of PRC2 activity is JARID2^8,13^, which is methylated at K116 and has a position −1 arginine (R115). Lastly, the natural H3 sequence also has a stabilizing arginine (R26) neighboring K27^23^.

Why might activation via automethylation be useful to PRC2? Automethylation is stimulated by histone H3 by ~2-fold *in vitro*, which suggests that the methylation loop serves to activate PRC2 in the presence of H3 tails. EZH2 automethylation also appears to activate PRC2 in response to SAM concentration, in the sense that increased SAM gives more automethylation, which then makes PRC2 more active. The RNA-binding sites of EZH2 include amino acids 489-494 in the methylation loop^24^, so it will be interesting to see if there is any cross-talk between RNA binding and automethylation.

There also remains an outstanding question of whether cellular methyltransferases and demethylases might be able to regulate PRC2 automethylation levels in order to modulate PRC2 function. We observed, quite interestingly, that a viral SET domain methyltransferase specific to H3K27 is capable of methylating PRC2 *in vitro* (data not shown). In addition to PRC2, another histone methyltransferase, G9a, has been demonstrated to automethylate itself. This automethylation provides wider substrate specificity and modulates binding of additional proteins^25,26^. In other examples of methylation of non-histone proteins, these marks act as important regulators of cellular signal transduction in MAPK and NF-κB signaling pathways ^27,28^. In these cases, crosstalk between histone and non-histone protein methylation also occurs and affects cellular processes such as chromatin remodeling, gene transcription, and protein synthesis.

What might be the therapeutic significance of understanding new PRC2 regulatory features? PRC2, one of the few enzymes in cells associated with gene silencing, is a natural candidate for epigenetic therapy. Indeed, cancers harboring mutations in the EZH2 subunit of PRC2 have been shown to be susceptible to small-molecule inhibitors that are currently in clinical development. For example, missense mutations in EZH2 are reported in follicular lymphoma and diffuse large B-cell lymphoma. The most prevalent mutation occurs at Y646 of EZH2, which is frequently altered to C, F, H, N, or S. These activating mutations cause H3K27 hypermethylation *in vitro* and *in vivo*, and they have been suggested to be associated with malignant transformation^29,30^. Early studies using highly-selective EZH2 inhibitors to treat follicular lymphoma and diffuse large B-cell lymphoma bearing these mutations have demonstrated some treatment success^31^. Based on our automethylation analysis, such EZH2 inhibitors should not only inhibit histone H3 methylation directly, but should also inhibit PRC2 activation through automethylation.

Intriguingly, the cancer genomic databases^32,33^ also report mutations in K510 and K515 of EZH2 (**Supplementary Figure 4**), residues that we described here to be key targets of automethylation and PRC2 autoregulation. The implication is that PRC2 might possibly be dysregulated at the level of the methylation loop in some cancers. Future *in vivo* studies are needed to test this hypothesis. Certainly, the data shown here provide new insights into the regulatory complexity of PRC2 and suggest that PRC2 evolved the ability to exquisitely fine-tune its activity in multiple ways. Our findings contribute to foundational knowledge for future studies pursuing an understanding of how PRC2 regulation can go awry in diseases.

## METHODS

### Protein expression and purification

Human PRC2-5m complexes, comprising EZH2, EED, SUZ12, RBBP4 and AEBP2 (UniProtDB entry isoform sequences Q15910-2, Q15022, O75530-1, Q09028-1, and Q6ZN18-1, respectively), were expressed in insect cells. In brief, standard Bac-to-Bac baculovirus expression system (Expression System) was used to generate baculovirus stocks based on standard protocol. Gp64 detection was used for tittering each baculovirus stock (Expression Systems). Sf9 cells (Invitrogen) were grown to a density of 2.0 x 10^6^ cells/ml, followed by infecting with equal amounts of baculovirus for each subunit. The cells were incubated for additional 72 h (27°C, 130 rpm), harvested and snap-frozen with liquid nitrogen for later purification.

A three-column purification scheme was used to purify PRC2 5-mer complexes as previously described ^9^. Briefly, insect cells were lysed in lysis buffer (10 mM Tris-HCl, pH 7.5 at 25 °C, 150 mM NaCl, 0.5% Nonidet P-40, 1mM TCEP) and cell lysate was bound to the amylose resin and washer thoroughly. The protein was eluted with 10mM maltose, followed by concentrating to ~15 mg/ml as final concentrations. PreScission protease was used to digest eluted protein at a mass ratio of 1:50 protease:protein. After overnight incubation at 4°C, cleavage efficiency was checked by SDS-PAGE. The cleaved protein was subject to 5 ml Hi-Trap Heparin column (GE, 17-0407-03) with a gradient over 35 column volumes from Buffer A (10 mM Tris-pH 7.5 at RT, 150 mM NaCl, and 1 mM TCEP) to Buffer B (10 mM Tris-pH 7.5 at RT, 2 M NaCl, and 1 mM TCEP), with a 1.5 ml/min flow rate. All the peak fractions were checked by SDS-PAGE and the PRC2 fractions were pooled and concentrated. The concentrated protein was subject to the final sizing column: HiPrep 16/60 Sephacryl S-400 HR with running buffer (250 mM NaCl, 10 mM Tris-HCl, pH 7.5 at RT, 1 mM TCEP-pH 7) with a flow rate of 0.5 ml/min. PRC2-peak fractions were checked with SDS-PAGE. The correct fractions were pooled and concentrated as above. Final protein concentration was calculated by nanodrop (UV absorbance at 280 nm). The ratio of absorbance at 260 nm/280 nm < 0.7 was observed, suggesting no nucleic acid contamination.

### *In vitro* histone methyltransferase assay

In each 10 µl reaction, recombinant PRC2-5m, H3 (NEB M2503S), and *S*-[methyl^14^C]-adenosylmethionine (PerkinElmer NEC363050UC) were mixed in methylation buffer (50 mM Tris-HCl pH 8.0 at 30°C, 100 mM KCl, 2.5 mM MgCl2, 0.1 mM ZnCl2, 2 mM 2-mercaptoethanol, 0.1 mg/ml bovine serum albumin, 5% v/v glycerol). All of the methylation reactions were incubated for 1 h at 30°C, followed by adding 4X loading dye to stop each reaction and heated at 95°C for 5 min. Each reaction was then loaded onto either 4-12% Bis-Tris gel (ThermoFisher NP0322BOX). Gel electrophoresis was carried out for 48 min at 180 V at room temperature. Gels were stained by InstantBlue for an hour and de-stained with water overnight. Three sheets of Whatman 3 mm chromatography paper were put underneath the gel and gels were scanned, followed by vacuum dried for 60 min at 80°C. Dried gels were subject to phosphorimaging plates and radioactive signal was acquired with a Typhoon Trio phosphorimager (GE Healthcare). Densitometry and analysis were carried out with ImageQuant software (GE Healthcare).

For the experiments of pre-incubating PRC2 with unlabeled SAM, 15 μM PRC2 was pre-incubated with 0.3 mM SAM first for 1 h at 30°C, followed by running through Quick Spin columns to remove all the cold SAM (Sigma-Aldrich 11273949001). The column was prepared and eluted according to manufacturer’s protocol. Then, the pure pre-incubated PRC2 was subjected to methylation assays as described above.

### Site-directed mutagenesis

Mutant EZH2 genes were generated using the QuickChange II site-directed mutagenesis kit (Stratagene). The appropriate mutations were confirmed by DNA sequencing.

### Mass spectrometry detection and analysis

Methylation experiments were set up as above. Mass spectrometry experiment and analysis were performed at the Core Facility of University of Colorado-Boulder. Samples were processed using standard protocol. In brief, protein samples (32 µg PRC2-5m complex) were diluted with an incubation buffer (50 mM Tris, pH 7.6, 5 mM CaCl2, 2 mM EDTA). 5 mM TCEP was used to reduce the reaction at 60 °C for 30 min, followed by alkylated with 15 mM iodoacetamide at room temperature for 20 min. 7.5 mM DTT was added to quench unreacted iodoacetamide. The reactions were digested with 0.6 µg of sequencing grade Arg-C (Promega) at 37 °C overnight, then desalted with Pierce C18 columns (Thermo Scientific) and dried with vacuum centrifugation. Prior to LC-MS/MS analysis, Buffer A (0.1% formic acid in water) was used to reconstitute the peptides.

For LC-MS/MS analysis, a Waters nanoACQUITY UPLC BEH C18 column (130 Å, 1.7 µm × 75 µm × 250 mm) was first equilibrated with 0.1% formic acid/3% acetonitrile/water, followed by peptide loading. Each load was an aliquot (5 µl, 1 µg) of the peptides. 0.1% formic acid/water was used as the mobile phase A and 0.1% formic acid/acetonitrile was used as phase B. The elution was done at the rate of 0.3 µl/min using gradients of 3 to 8% B (0-5 min) and 8 to 32% B (5-123 min). A LTQ Orbitrap Velos mass spectrometer was used for MS/MS. The precursor ions were scanned between 300 and 1800 m/z (1 × 10^6^ ions, 60,000 resolution). The 10 most intense ions were selected with 180 s dynamic exclusion, 10 ppm exclusion width, repeat count = 1, and 30 s repeat duration. Ions were excluded based on unassigned charge state and MH+1 from the MS/MS. 500 ms for FT (one microscan) and 250 ms for LTQ were set as maximal ion injection times. The automatic gain control was 1 × 10^4^ and the normalized collision energy was set as 35% with activation Q 0.25 for 10 ms.

For database search, MaxQuant/Andromeda (version 1.5.2.8) was used. The raw files from LTQ-orbitrap were processed. The peak was searched against Uniprot human proteome. In the search, Arg-C specificity with a maximum of two missed-cleavages was used. Several modifications, including carbamidomethyl modification on cysteine as a fixed modification and protein N-terminal acetylation, oxidation on methionine, and methylation on lysine or arginine as variable modifications, were set. In addition, search tolerance was set as 4.5 ppm main search tolerance for precursor ions and match tolerance was placed as 0.5 Da MS/MS match tolerance, searching top 8 peaks per 100 Da. Finally, false discovery rate was put as 0.01 with minimum seven amino acid peptide length.

### CRISPR-editing of HEK293T cells

Two plasmids were made for the CRISPR-editing. A CRISPR plasmid encoding Cas9 and the guide RNA was made by inserting the sgRNA sequence (CAGACGAGCTGATGAAGTAA) targeting exon 2 of the *EZH2* gene in pX330 as previously described^34^. Two donor plasmids carrying either the WT or mutant *EZH2* cDNA were made by assembling the following fragments into a previously described donor plasmid^35^: left homology arm (-951 to −14, relative to the ATG start codon), *EZH2* cDNA, *EZH2* 3’UTR (872 bp immediately after the stop codon), 3X SV40 polyadenylation sites, 1X bGH polyadenylation site, SV40 promoter, puromycin resistance ORF, T2A self-cleavage site, mCherry ORF, SV40 polyadenylation site and right homology arm (+25 to +830, relative to ATG). 1.2 µg of CRISPR plasmid and an equal amount of donor plasmid were transfected to 1 million HEK293T cells in a 6-well plate using Lipofectamine 2000 according to the manufacturer’s instructions. Cells were passaged to a 15-cm plate after one day and 1 µg/mL of puromycin was added to the culture two days later. Cells were selected in the presence of puromycin for one week and the surviving cells were pooled into a well of a 6-well plate. A Cre-GFP plasmid was transfected and cells with both GFP and mCherry signal were selected and sorted into 96-well plates using flow cytometry after 24 hours. When each clone reached confluency, cells were passaged and a fraction was used for genomic DNA extraction as previously described^36^. Four primers were used for verification of the correct genome editing: P1 gctgcagcatcatctaacctgg, P2 cagtgagtcagaaaaccttgctc, P3 atcatctcggtgatcctccag and P4 tgagcagtcctgaaagcagttatt. PCR products were analyzed on a 1% agarose TAE gel.

### Western Blot

CRISPR-edited HEK293T cells were harvested and 1x NuPAGE LDS Sample Buffer (Life Technologies) supplemented with Benzonase nuclease (Novagen) to 10 U/μl was used for preparing samples for SDS-PAGE. Standard Western Blot protocol was used and antibodies for detections include: H3K27me1 (Active Motif, 61065, 1:1000), H3K27me2 (Cell Signaling, 9278S, 1:1000), H3K27me3 (Cell Signaling 9733S, 1:500), EZH2 (Cell Signaling 5246S, 1:1000) and beta-actin (ThermoFisher, MA1-91399, 1:10000). Validation of each antibody can be found on the manufacturers’ websites.

## ACKNOWLEDGEMENTS

We thank members of the Cech lab for useful discussion. We are grateful to Jeremy Balsbaugh from the core facility at University of Colorado-Boulder for performing the initial Mass Spectrometry experiments. We thank Chen Davidovich (Monash University, Australia) for the initial discussion of this project. T.R.C. is an investigator of the Howard Hughes Medical Institute.

---

**Supplementary Figure 1.**
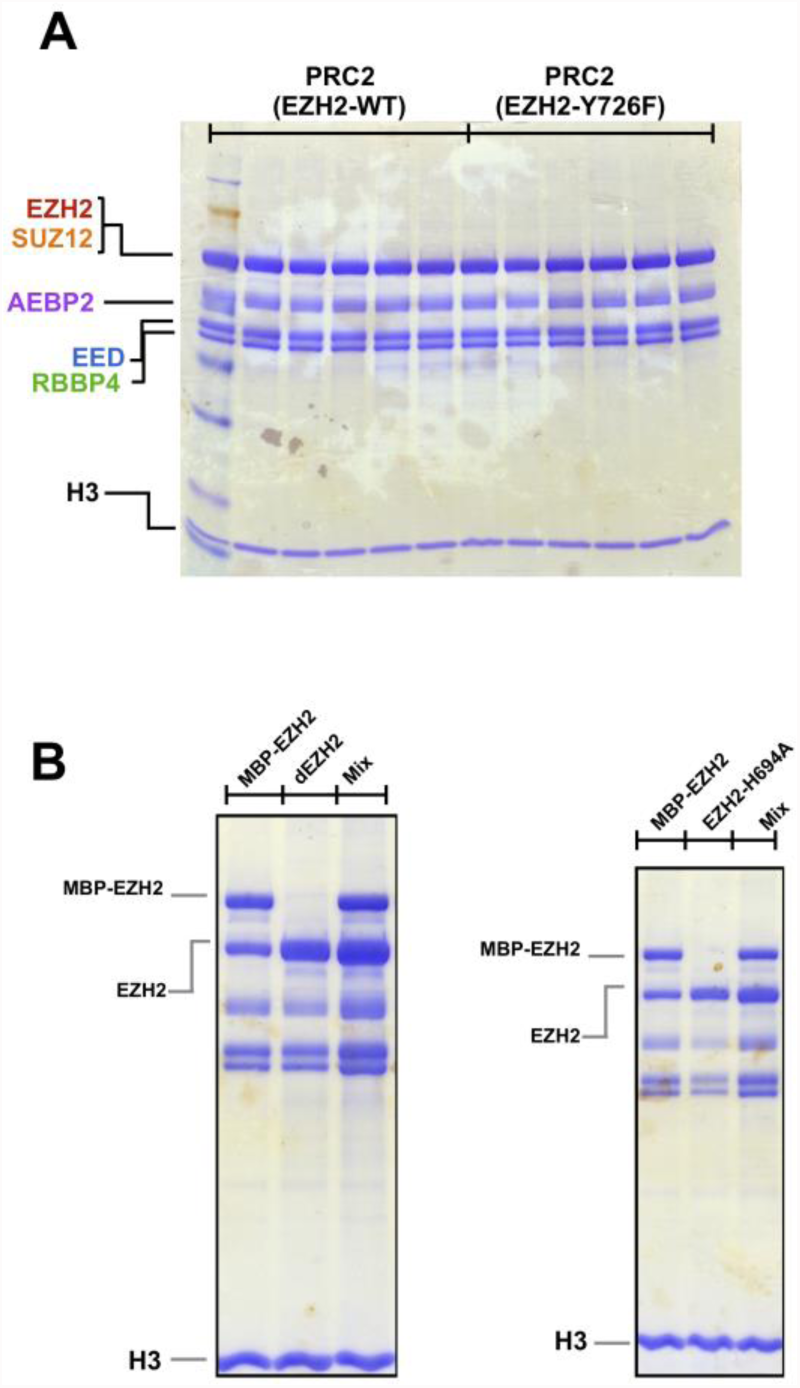
Coomassie-stained gels of reactions of PRC2 and H3. **A.** Coomassie stain of Figure 4B. **B.** Coomassie stain of Figure 4D.

**Supplementary Figure 2.**
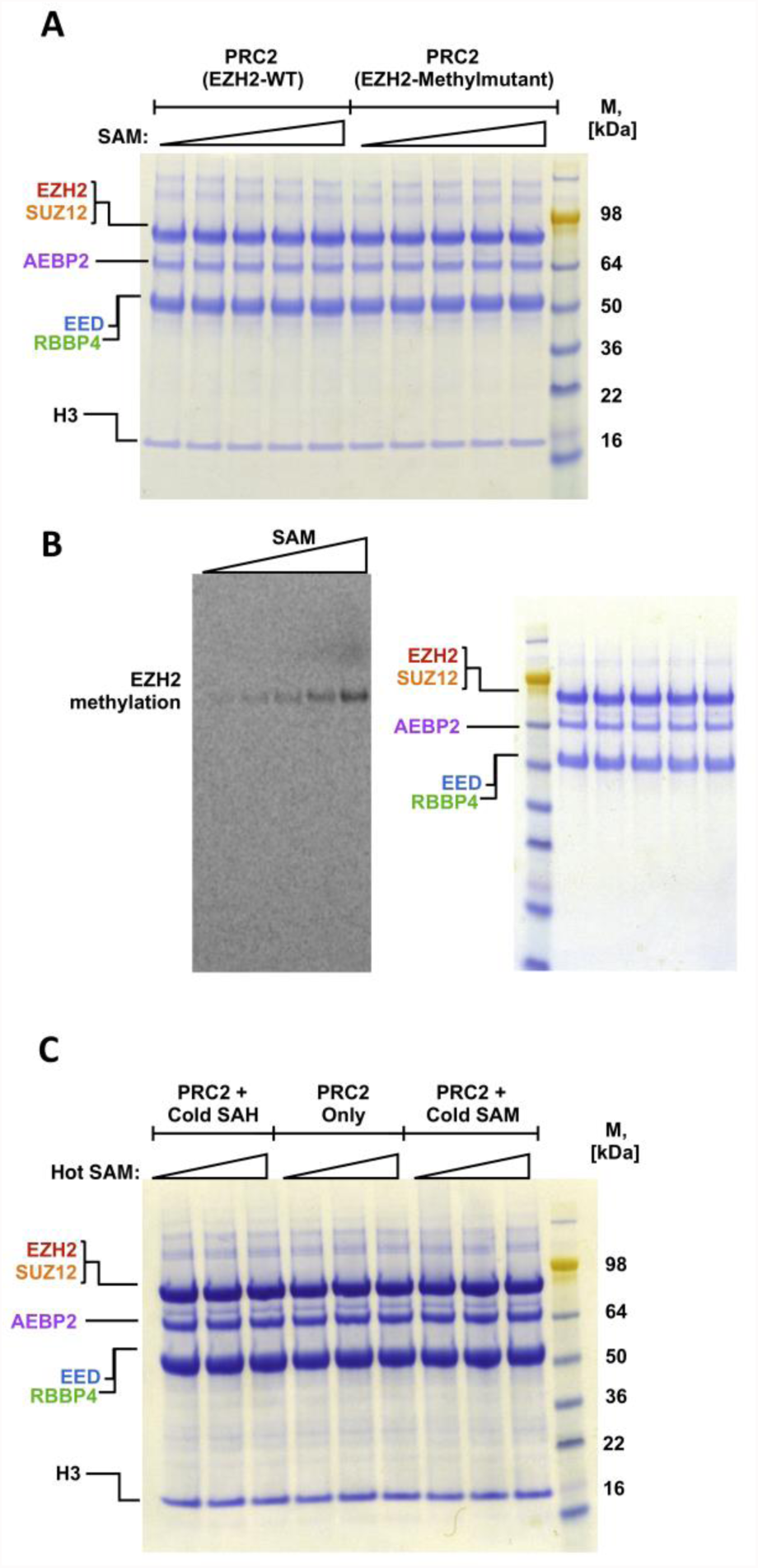
Automethylation of PRC2 activates histone substrate methylation. **A.** Coomassie stain of Figure 5A. **B.** Left, HMTase assays showing EZH2 is automethyated with SAM. HMTase assays were performed in the presence of cofactor ^14^C-SAM; Right, Coomassie stain of the gel shown in the left. **C.** Coomassie stain of Figure 5D.

**Supplementary Figure 3.**
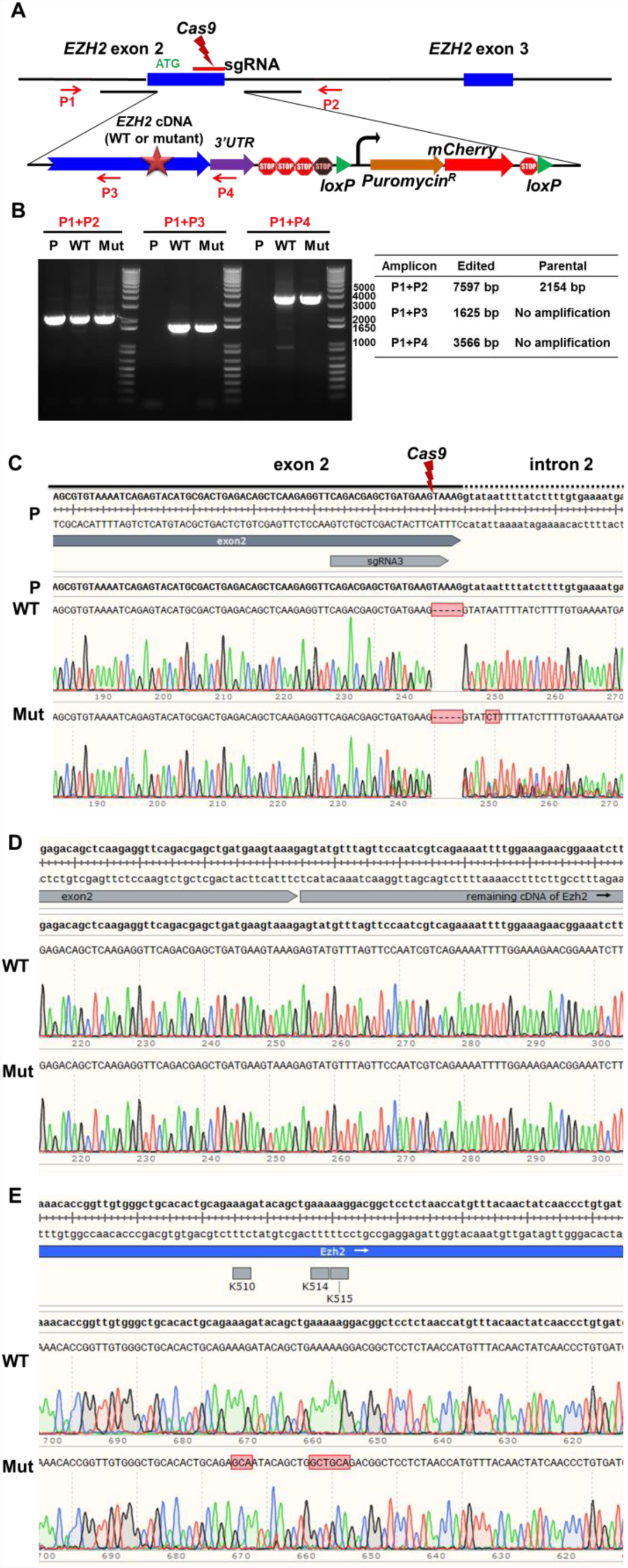
CRISPR editing of *EZH2* gene to generate methylmutant alleles. **A**. Schematic representation of the CRISPR-Cas9 genome-editing strategy. Binding sites of the four primers (P1-4) used for PCR validation of the correct integration are indicated by the red arrows. **B.** PCR products were generated from genomic DNA from HEK293T single clones (WT and Mut) and unedited parental cells (P). The expected sizes of PCR products for each cell line are indicated in the table to the right of the gel. The sizes of the P1+P2 PCR products for both WT and Mut are similar to that of the parental line, indicating that both edited strains are heterozygous. Absence of an upper band representing the edited allele was caused by the PCR bias towards the shorter product (the unedited allele). **C.** Sanger sequencing analysis of the P1+P2 PCR product of WT and Mut indicates that five nucleotides are deleted at the Cas9 cleavage site, resulting in a frame shift and early termination for the unedited allele. This deletion was observed in both the WT and Mut strain, and “P” indicates the parental sequences. **D.** Sequencing results of the P1+P4 PCR product of WT and Mut near the end of the exon 2 indicate correct integration of the cDNA at the Cas9 cleavage site for both strains. E. Sequencing results of the P1 + P4 PCR product of WT and Mut near the K510, K514 and K515 residues indicate the alanine mutations only exist in the Mut strain but not the WT strain.

**Supplementary Figure 4.**
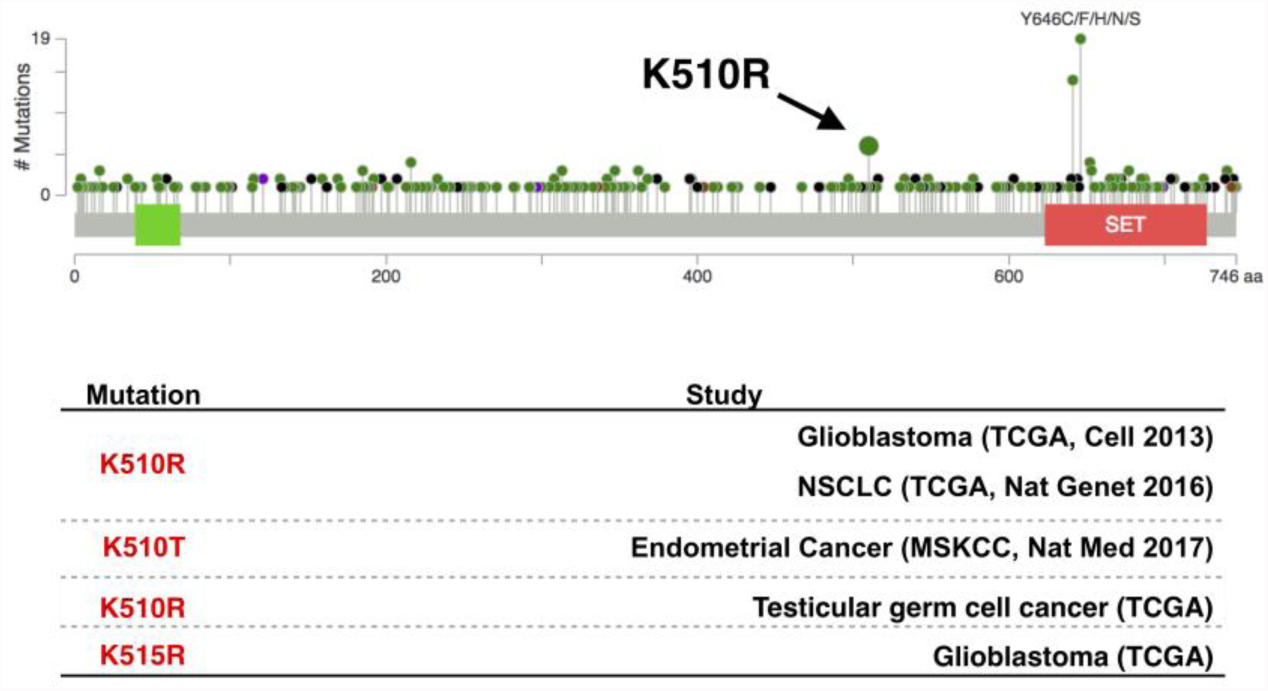
Spectrum of EZH2 mutations in cancer genomics database. **Top,** Frequent mutations occur at EZH2 site Y646. Mutations at sites K510 and K515 are also observed (K510 indicated in the visualization by the black arrow). Data visualized using cBioPortal. **Bottom,** Mutations in the methylation residues matched with the respective study.

